# Genetic dissection of the pluripotent proteome through multi-omics data integration

**DOI:** 10.1101/2022.04.22.489216

**Authors:** Selcan Aydin, Duy T. Pham, Tian Zhang, Gregory R. Keele, Daniel A. Skelly, Matthew Pankratz, Ted Choi, Steven P. Gygi, Laura G. Reinholdt, Christopher L. Baker, Gary A. Churchill, Steven C. Munger

## Abstract

Genetic background is a major driver of phenotypic variability in pluripotent stem cells (PSCs). Most studies of variation in PSCs have relied on transcript abundance as the primary molecular readout of cell state. However, little is known about how proteins, the primary functional units in the cell, vary across genetically diverse PSCs, how protein abundance relates to variation in other cell characteristics, and how genetic background confers these effects. Here we present a comprehensive genetic study characterizing the pluripotent proteome of 190 unique mouse embryonic stem cell lines (mESCs) derived from genetically heterogeneous Diversity Outbred (DO) mice. The quantitative proteome is highly variable across DO mESCs, and we identified differentially activated pluripotency-associated pathways in the proteomics data that were not evident in transcriptome data from the same cell lines. Comparisons of protein abundance to transcript levels and chromatin accessibility show broad co-variation across molecular layers and variable correlation across samples, with some lines showing high and others low correlation between these multi-omics datasets. Integration of these three molecular data types using multi-omics factor analysis revealed shared and unique drivers of quantitative variation in pluripotency-associated pathways. QTL mapping localized the genetic drivers of this quantitative variation to a number of genomic hotspots, and demonstrated that multi-omics data integration consolidates the influence of genetic signals shared across molecular traits to increase QTL detection power and overcome the limitations inherent in mapping individual molecular features. This study reveals transcriptional and post-transcriptional mechanisms and genetic interactions that underlie quantitative variability in the pluripotent proteome, and in so doing provides a regulatory map for mouse ESCs that can provide a rational basis for future mechanistic studies, including studies of human PSCs.

## Introduction

Pluripotent stem cells (PSCs) hold great potential for modeling human disease and advancing regenerative medicine (Hamazaki et al., 2017), but variation in the derivation, stability, and differentiation of individual cell lines impedes progress toward these goals (Ortmann and Vallier, 2017; Volpato and Webber, 2020). Genetic background contributes significantly to phenotypic variation in human and mouse PSCs (Czechanski et al., 2014; Ortmann and Vallier, 2017). Systems genetics experiments can identify the loci that harbor genetic variants, and can associate phenotypic variability with regulatory networks that are affected by these variants (Byers et al., 2022; Carcamo-Orive et al., 2017; Kilpinen et al., 2017; Mirauta et al., 2020; Panopoulos et al., 2017; Skelly et al., 2020).

Most studies addressing phenotypic variability in PSCs have focused on transcriptional regulation using measures of chromatin state and transcript abundance, due in part to the relatively low cost of RNA and DNA sequencing. However, cellular phenotypes are largely determined by proteins, and the effects from genetic variation on chromatin states and transcripts may be buffered, amplified, or even reversed by post-transcriptional processes acting on protein abundance (Chick et al., 2016; Mirauta et al., 2020). Previous studies in cell and animal models have found a surprising level of disagreement between protein and transcript abundance (Buccitelli and Selbach, 2020; Gygi et al., 1999; Maier et al., 2009; Vogel and Marcotte, 2012); this high discordance was also observed in differentiating mouse embryonic stem cells (mESCs; (van den Berg et al., 2017)). Genetic analyses suggest that stoichiometric buffering acting on protein complexes may attenuate transcriptional variation of complex-forming proteins in adult mouse tissues (Chick et al., 2016; Keele et al., 2021), and translational output was recently shown to provide strong feedback on chromatin state and transcription to drive self-renewal in mESCs (Bulut-Karslioglu et al., 2018). These findings suggest that post-transcriptional regulation of protein abundance may play a significant role in pluripotency maintenance and differentiation in PSCs.

We previously derived a panel of mESCs from Diversity Outbred mice (DO mESCs). The DO mice are an outbred population derived from eight inbred founder strains with high genetic diversity, and a population structure that is optimized for genetic mapping and causal variant discovery (Churchill et al., 2012; Skelly et al., 2020). We maintained DO mESCs in sensitized culture conditions to amplify genetic differences in the pluripotent ground state and analyzed transcriptome and chromatin state data to map genetic modifiers underlying this variability (Skelly et al., 2020). We demonstrated that genetic variation influences chromatin accessibility and transcript abundance to alter the stability of the ground state, as measured by markers of pluripotency and capacity for selfrenewal (Skelly et al., 2020). We showed that genetic background can bias differentiation propensity of mESCs through its effects on Wnt signaling activity (Ortmann et al., 2020). These studies demonstrated the power of this resource for discovery of genetic drivers and molecular mechanisms that underlie variation in the maintenance of the pluripotent ground state and differentiation propensity of mESCs.

In the current study we expand on the previous work by investigating how genetic effects are mediated by the proteome. We quantified proteins by multiplexed mass spectrometry across the same panel of DO mESC lines. As with our previous analysis of chromatin accessibility and transcript abundance, we find the quantitative proteome to be highly variable across these cell lines. Genetic mapping analysis identified significant protein quantitative trait loci (pQTL) for 20% of all measured proteins. One third of the pQTL appear to uniquely affect protein abundance independently from transcript levels – presumably through post-transcriptional mechanisms. Thus, these signatures of genetic effects on proteins were not detected in our earlier analysis of transcript abundance. The remaining pQTL colocalize with previously identified QTL for transcript abundance (eQTL) and/or chromatin accessibility (caQTL), consistent with transcriptional regulation. We applied multi-omics factor analysis (MOFA) to identify latent factors that account for the variability in gene regulatory signatures across these three layers of molecular data (Argelaguet et al., 2018). Genetic mapping of the latent factors identified previously reported QTL “hotspots” as well as novel regulatory loci. We show that multi-omics integration increases power to detect genetic drivers of broad regulatory signatures compared to QTL mapping of individual molecular traits. We further show how genetic variation affects transcriptional and post-transcriptional gene regulation to drive variation in ground state pluripotency. The resulting regulatory map for mouse ESCs can provide a rational basis for future mechanistic studies, including studies of human PSCs.

## Results

### The pluripotent proteome of genetically diverse mESCs

We applied multiplexed mass spectrometry to quantify relative protein abundance in 190 unique DO mESC lines (Fig 1A). In total, we detect 7,432 proteins in at least half and 4,794 proteins in all the cell lines. The list of proteins detected in mESCs is overrepresented for those involved in cellular metabolism (e.g., organic acid metabolic process), post-transcriptional processes (e.g., translation, mRNA processing), and protein complexes (e.g., spliceosome, proteasome). By contrast, transmembrane proteins and transcription factors are overrepresented among the genes showing expression in the RNA-seq data but not detected in the proteomics data (Table S1). Transmembrane proteins contain both hydrophilic and hydrophobic subunits making them less soluble (Schey et al., 2013) and therefore harder to isolate in untargeted proteomics analysis. In addition, the probability of detecting an individual protein is dependent on its transcript abundance (Fig S1A); expressed genes at the lower threshold for transcript abundance (average count = 1) have a protein detection rate of less than 60% (Fig 1B). This includes transcription factors, which as a group exhibit lower mean transcript abundance, presumably resulting in lower levels of detectable protein (Fig S1B). By contrast, proteins encoded by genes with high transcript expression—a group that includes many ribosomal and mitochondrial proteins—are detected at a much higher rate (>90%) (Fig 1B). Of note, ribosomal genes segregate high genetic diversity in the DO which makes short read alignment to them particularly challenging, and the observed dip in protein detection rate for the highest expressed genes is likely due to read alignment errors causing inflated transcript abundance estimates for some ribosomal genes.

**Figure 1.**
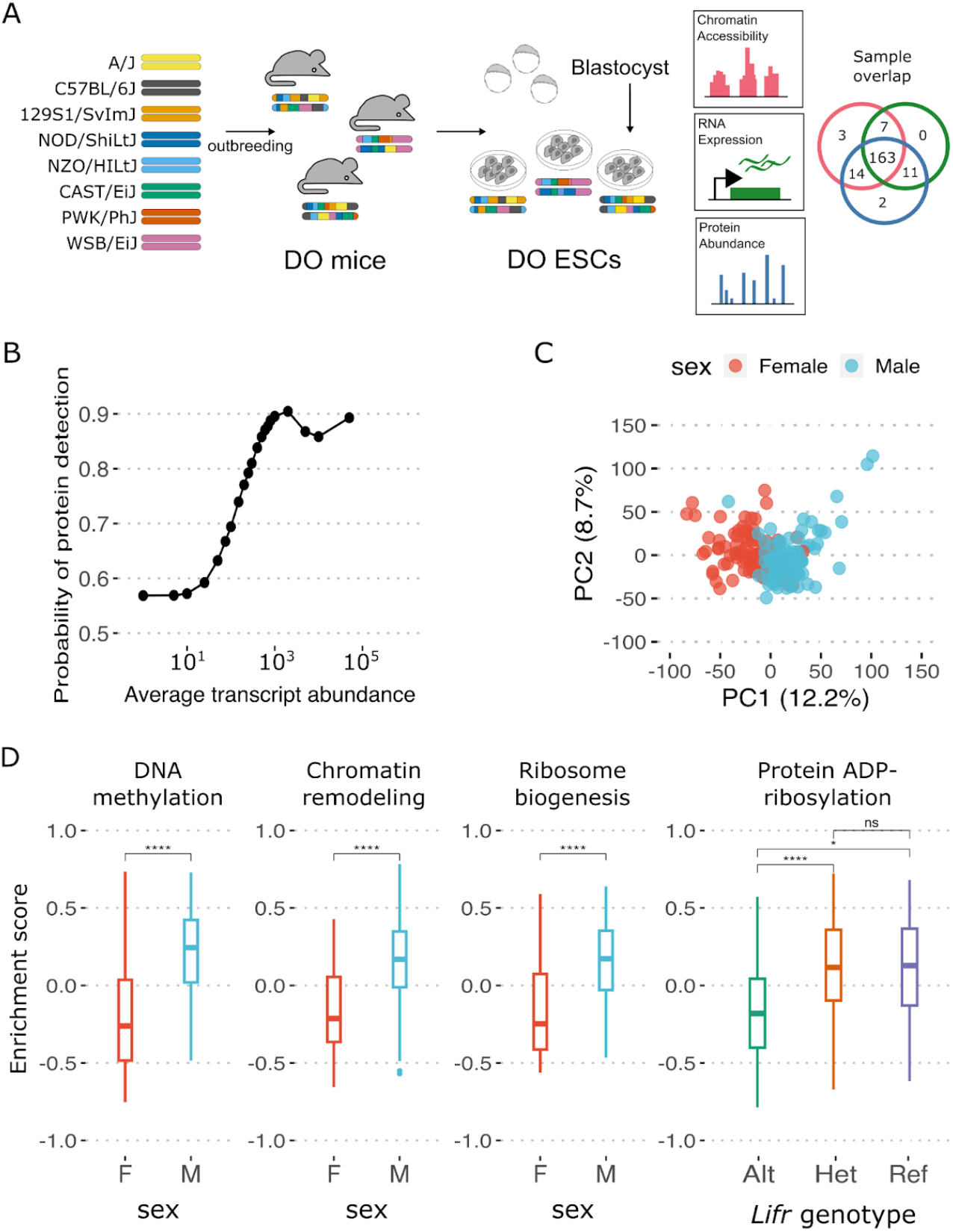
Overview of the quantitative proteome in genetically diverse mESC lines. (A) Nearly 200 embryonic stem cell lines were established from blastocysts of Diversity Outbred mice, and quantified using ATAC-seq, RNA-seq, and multiplexed mass spectrometry; 163 lines have all three measures. (B) Protein detection rate is linked to transcript abundance. The probability of a gene to have protein abundance measurement given its average transcript abundance among 174 mESCs with both transcriptome and proteome data. (C) Principal component analysis reveals sex as a significant source of variation among DO mESC proteomes. PC1 and PC2 for 190 mESCs are plotted and colored by sex. (D) Enrichment scores obtained from GSVA for Gene Ontology Biological Processes (GO:BP) showing significant differences between mESCs with different sexes and genotypes at the *Lifr* locus are plotted. GO:BP DNA methylation, chromatin remodeling and ribosome biogenesis show significantly higher enrichment in males in comparison to females and, protein ADP-ribosylation shows significantly higher enrichment in mESCs with at least one copy of the reference allele in comparison to ones carrying two copies of the alternative allele at the *Lifr* locus (two-way anova followed by Tukey’s HSD, *: p value < 0.05, ****: p value < 0.00005).

The mESC proteome is highly variable across cell lines (Fig S1D, E), and the highest sample-level correlations are observed between replicate lines and those derived from related individuals, consistent with genetic effects. Principal Component Analysis (PCA) points to chromosomal sex as the largest component of variance across samples (12.2%, Fig 1C), and sexually dimorphic expression of X-linked proteins drives most of this variation. This expression bias is likely due to gene dosage effects from two active X chromosomes in XX mESCS (Epstein et al., 1978; Kratzer and Gartler, 1978). Overall, more than half of all proteins expressed in DO mESCs exhibit variable expression linked to sex (n = 4,106 / 7,432, p < 0.05), including pluripotency factors SOX2, ESRRB, KLF2, KLF4, SALL4, UTF1, NR5A2 and LIN28A (Kalkan et al., 2017). To identify pathways and biological processes that vary across cell lines, we performed gene set variation analysis (GSVA) (Hänzelmann et al., 2013). GSVA is an unsupervised method that estimates variation in the activity (expression) of annotated gene sets across a population. The resulting enrichment scores can be used to compare groups of samples, or as traits themselves for downstream analyses and genetic mapping. For example, female and male samples vary in many cellular processes and protein complexes (Table S2); male cell lines show higher enrichment for DNA methylation, histone modification and chromatin remodeling in comparison to female cell lines, consistent with previous studies (Schulz et al., 2014; Werner et al., 2017) (Fig 1D). We find that ribosome biogenesis genes are overrepresented in weightings of the first principal component, with higher abundance in male cell lines compared to female lines (p < 5 x 10-^4^) (Fig 1D). Male cell lines also showed higher abundance of proteins in complexes involved in chromatin remodeling and ribosome biogenesis (p < 5 x 10^-4^), consistent with findings from other proteome analyses (Keele et al., 2021; Romanov et al., 2019). In addition, the proteomes of cell lines with different genotypes at the *Lifr* locus, where those with the reference allele have increased stability in the pluripotent state (Skelly et al., 2020), differ significantly in several biological processes (Table S2). For example, cell lines with at least one copy of the reference *Lifr* allele showed higher abundance of proteins with regulatory roles in ADP-ribosylation, the transfer of ADP-ribose moieties (derived from NAD) to protein amino acids (Fig 1D). This post-translational histone modification is catalyzed by poly-ADP-ribose polymerases, two of which (PARP1/ARTD1, PARP7/TIPARP) have been shown in mESCs to occupy and maintain an active epigenetic state at key naïve pluripotency genes including *Nanog, Oct4/Pou5f1, Sox2*, and *Rex1/Zfp42* (Roper et al., 2014). GSVA enrichment of ADP-ribosylation proteins as well as several other pluripotency and differentiation pathways are observed only in the proteomics data and not evident in the transcriptome GSVA results (pathways highlighted in Table S2).

Extracellular proteins are overrepresented among the most variable proteins, while members of protein complexes are overrepresented among the least variable proteins. These include chaperonin containing T (CCT) complex; minichromosome maintenance (MCM) complex; proteasome; and spliceosome (false discovery rate [FDR] < 0.05). The overrepresentation of these functional groups of proteins is not simply due to having low or high abundance or lacking variability (Fig S1F). Targets of the transcription factor REX1, a marker of naïve pluripotency (Kalkan et al., 2017), are also overrepresented among the least variable proteins, despite REX1 protein itself being highly variable across ESCs (Fig S1G). REX1 is known to act as a repressor (Guallar et al., 2012), and the lowest REX1-expressing mESC lines may still exceed a threshold required to efficiently repress its target genes. Alternatively, the effects of variable REX1 protein abundance on downstream targets may be buffered. Reports in the literature indicate that REX1 may be dispensable for pluripotency maintenance (Masui et al., 2008), which would support the existence of compensatory mechanisms.

To further characterize the influence of physical interactions on protein abundance, we looked at the covariation among complex-forming proteins. Members of protein complexes have a lower median coefficient of variation (CV) than non-complex forming proteins, suggesting that physical interactions on the whole act to dampen the variation in protein abundance (Fig S2A). We observed that complex-forming proteins tended to covary in their abundance more than proteins that are not known to physically interact (Fig S2B), in line with previous studies (Romanov et al., 2019). We quantified this covariation, or “cohesiveness”, of complexes using median pairwise Pearson correlation between complex members; higher median correlation indicates tighter co-regulation of subunits and a stable maintenance of stoichiometry. We ranked protein complexes by their cohesiveness and observed that the most stable 10% of complexes (n = 17) were associated with the cell cycle, protein modification, and translation machinery—consistent with our analysis of individual proteins and published proteome studies of human and mouse cell lines and tissues (Hansson and Krijgsveld, 2013; Ori et al., 2016; Romanov et al., 2019) (Fig 2). Several complexes involved in protein trafficking and transcriptional regulation are also highly cohesive. By comparison, the least cohesive 10% of complexes is enriched for those associated with chromatin remodeling. Sex differences in complex cohesiveness are also observed; for example, protein constituents of the cytoplasmic small and large ribosomal subunits, the mitochondrial small ribosomal subunit, and HOPS complex are significantly more cohesive in XY than XX mESCs (p < 5 x 10-^4^) (Fig S2C). The molecular basis of these sex differences remains to be established; however, they are also observed in adult mouse liver (Keele et al., 2021) and heart proteomes (Gyuricza et al., 2022), and are therefore unlikely to play a unique role in establishing or maintaining pluripotency.

**Figure 2.**
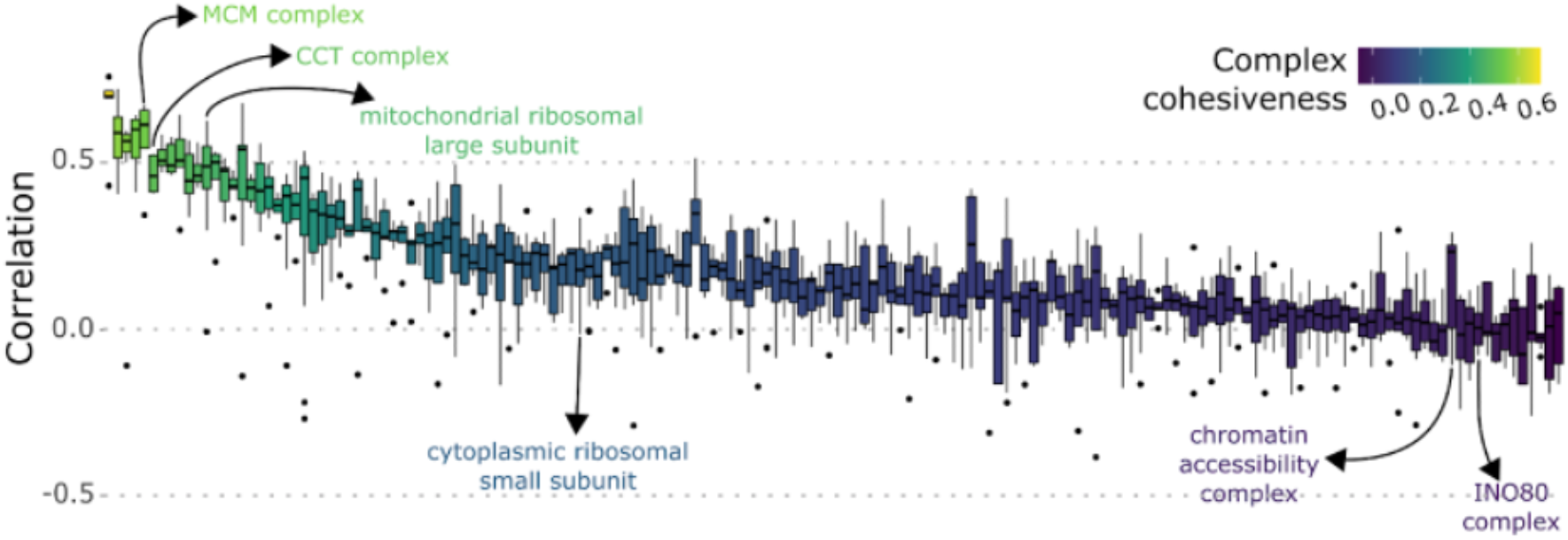
Protein subunit cohesiveness varies considerably among 164 complexes in DO mESC lines. For each complex, pairwise Pearson correlations were calculated between all protein subunits and summarized as a boxplot. Boxplots are ordered and colored based on their median pairwise correlation, with more cohesive complexes on the left. Specific examples of the stable (most cohesive 10%) and variable (least cohesive 10%) complexes are highlighted.

Cohesiveness is associated with the fidelity of a complex’s constituent proteins. Complexes tend to be more cohesive if they consist of protein subunits that contribute to no or few other complexes (n ≤ 3) and less cohesive if they contain more promiscuous protein subunits that contribute to many other complexes (up to 10) (Fig S2D). We identified 51 proteins that each contribute to at least 3 distinct complexes, and we assessed their covariation with members of each complex. Some of these promiscuous proteins show no preference among their annotated complexes, as evidenced by similar co-abundance to members across all complexes, while others show preferential membership to—or strict avoidance of—specific complexes in DO mESCs (Fig S2E). Examples of each include: CDK8, which shows high agreement with subunits of all three complexes it belongs to (median Pearson correlation coefficient [*r*] = 0.2), and HDAC3, which only shows high agreement with the HDAC3/NCOR complex (median *r* = 0.2) but not the Rpd3L (median *r* = −0.06), Emerin C32 (median *r* = −0.07), or Emerin regulatory complexes (median *r* = −0.03). Beta Actin (ACTB) isoforms appear to prefer different complexes in mESCs, with ACTB-208 showing higher agreement to members of the P2X7 receptor signaling and emerin complexes (median *r* = 0.2), and ACTB-201 showing higher agreement to members of the BAF complex (median *r* = 0.2). These findings of differential complex cohesiveness cannot be seen in the transcript data. Further studies are needed to elucidate the molecular mechanisms underlying promiscuous complex members’ preferences and their potential functional impacts on stability of the ground state.

### Protein abundance co-varies with chromatin accessibility and transcript abundance

The pluripotent state is established and maintained by a gene regulatory cascade that orchestrates changes across multiple molecular layers from chromatin accessibility to transcript and protein abundance (Nichols and Smith, 2009). To better understand these multi-layered regulatory interactions and to identify proteins with potentially important roles, we looked at the covariation of proteins with measures of chromatin accessibility (ATAC-seq) and transcript abundance (RNA-seq) across our panel of genetically diverse DO mESCs.

We compared protein abundance (n = 7,148) to chromatin accessibility (n = 99,159 peaks) and found that many proteins were most highly correlated with chromatin accessibility in the region proximal to their protein-encoding gene (Fig S3A), consistent with our earlier observation of high concordance between transcript abundance and local open chromatin (Skelly et al., 2020) and likely indicative of transcriptional regulation. We identified 37 proteins whose abundance co-varied with chromatin accessibility genome-wide at 100 or more ATAC-seq peaks (abs(*r*) > 0.5; evident as horizontal bands in Fig S3A), which suggests that these proteins may play a role in chromatin remodeling and gene regulation. The list includes both well-characterized pluripotency regulators and proteins with no previously reported role in pluripotency maintenance (Table S3). For example, the abundance of ID1, a transcription factor critical in the maintenance of ES cell self-renewal and regulation of lineage commitment (Romero-Lanman et al., 2012), covaries significantly (positively and negatively) with chromatin accessibility at 112 ATAC-seq peaks across the genome (Fig 3A). Other proteins with potential roles in pluripotency maintenance include: AHDC1, a putative DNA-binding protein previously shown to physically interact with TCF7L1 (TCF3), a transcription factor involved in pluripotency regulation (Moreira et al., 2018; Wray et al., 2011); and UHRF2, a ubiquitin ligase identified as a target of epigenetic control during self-renewal (Walker et al., 2010). For almost half of these proteins, we find that their covarying ATAC-seq peaks are overrepresented in binding sites active in ESCs for TRP53 (n = 28) and naïve pluripotency factors NANOG, ESRRB, and PRDM14 (n = 19, 18, 16 at FDR < 0.05) (Kalkan et al., 2017). Many of these covarying chromatin peaks are proximal to genes involved in cellular response to leukemia inhibitory factor (LIF), providing further evidence for their roles in establishing and/or maintaining pluripotency (FDR < 0.05). Notably, only six of the 37 genes covary with chromatin accessibility for both protein and transcript abundance, while 29 exhibit protein-specific correlations to chromatin, consistent with post-transcriptional regulation of these chromatin modifying proteins (Table S3).

**Figure 3:**
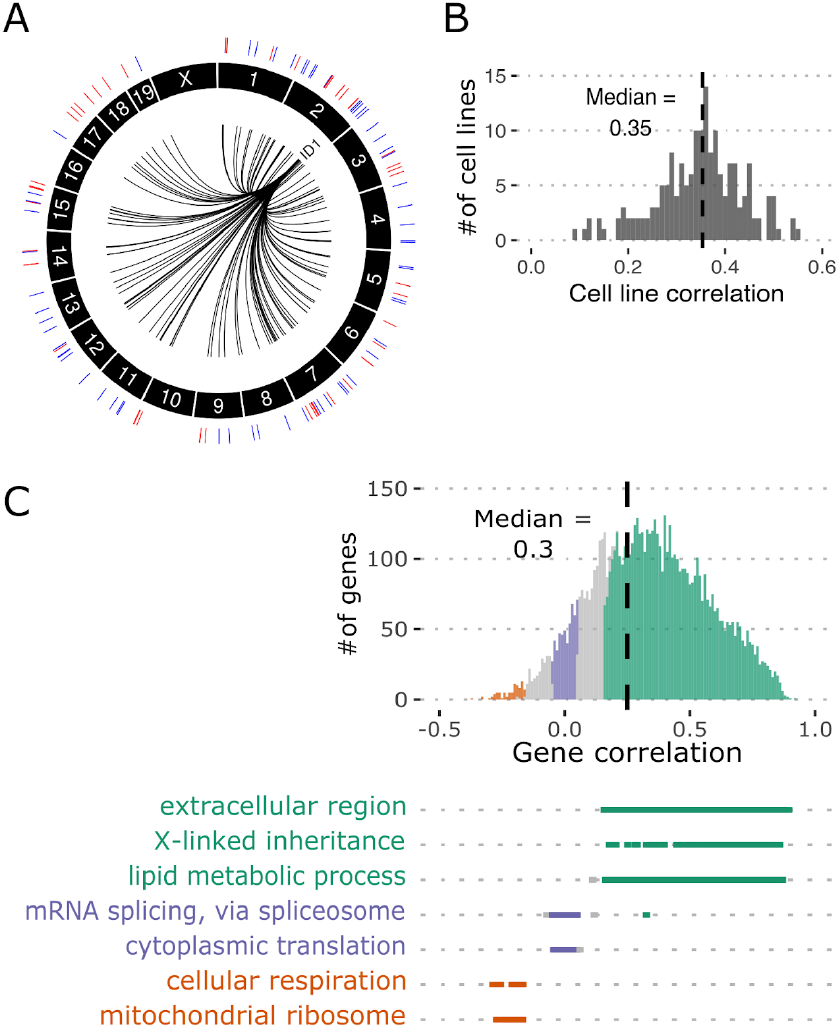
The quantitative proteome co-varies with chromatin accessibility and the transcriptome. (A) ID1 protein abundance shows high correlation to many chromatin regions across the genome. Circos plot showing ATAC-seq peaks where chromatin accessibility is positively (red) and negatively (blue) correlated with ID1 protein abundance (n = 112, abs(correlation) >0.5). (B) DO mESCs show a wide range of correlations between their transcriptome and proteome. Histogram of Pearson correlation coefficients between the transcriptome and proteome of DO mESC cell lines with matching genotypes (n = 174). (C) Genes show variable agreement between transcript and protein abundance in DO mESCs. Histogram depicting the distribution of pairwise Pearson correlation coefficients between transcript and protein abundance of genes with characteristic GO terms overrepresented in each category annotated underneath in matching colors (green: significantly positively correlated, orange: significantly negatively correlated, purple: genes with little or no correlation).

We next examined the concordance between protein and transcript abundance in DO mESC lines. For genes where we detect both (n = 7,241), protein and transcript abundance are broadly positively correlated in their magnitude and variance (*r* = 0.5, p < 2.2e-16, Fig S3B, C). Similar studies in human iPSCs found that many proteins that varied in abundance did not show variation in their cognate RNAs (Mirauta et al., 2020). We see a similar trend for a small number of proteins (n = 180) where protein abundance is highly variable across cell lines without similar variation at the transcript level. On the other extreme, genes with high variation in transcript abundance but lacking variation at the protein level (n = 111) were overrepresented for ribosomal proteins. Surprisingly, the overall agreement between protein and transcript levels within a cell line appears to vary considerably across the DO mESCs (*r* range 0.1 - 0.6) (Fig 3B). We ruled out sample mix-ups as a potential reason for the low concordance in some cell lines (Fig 3B), and even the lowest observed sample correlation is still well above a null distribution of correlation values from permuted sample assignments (Fig S3D). Looking at individual genes across DO mESC lines, we see a wide range of variation in the correlation between protein and transcript levels, where many are highly positively correlated while others are negatively correlated (Fig 3C). The larger group of genes showing positive transcript-protein correlation (n = 5,530, *r* > 0.16, p < 0.05) are over-represented for proteins involved in X-linked inheritance, lipid metabolism, and membrane proteins (Fig 3C). The smaller group of genes with significantly negatively correlated transcript and protein levels (n = 82, r < −0.16, p <0.05) are enriched for those with roles in cellular respiration and mitochondrial translation (Fig 3C). The list of genes exhibiting little or no correlation in their transcript and protein levels (abs(*r*) < 0.05, n = 498) are enriched for functions associated with mRNA splicing and cytoplasmic translation (Fig 3C). Stronger correlation of transcript and protein abundance is seen for genes that do not form protein complexes (Fig S3E), further supporting the idea that complexes place physical constraints on protein abundance that can serve to buffer against transcriptional variation (Chick et al., 2016; Keele et al., 2021).

### Genetic characterization of the pluripotent proteome

Variation in protein abundance across DO mESC lines appears to be driven at least in part by genetic background, with more than 90% of measured proteins estimated to have non-zero heritability (median *h^2^* = 0.25). To identify specific genomic loci underlying this quantitative variation in individual proteins, we performed protein quantitative trait locus (pQTL) mapping. For over 20% of expressed proteins (n = 1,555 / 7,432) we detected one or more pQTL, with a total of 1,677 pQTL (LOD > 7.5, permutation genome-wide p < 0.05, corresponding to an FDR = 0.058) (Fig 4A). Of these, nearly two-thirds (n = 1,056) are local and map to within ±5 Mb of the corresponding protein-encoding gene. We found many fewer distant pQTL (n = 621) that map outside of the local genomic window. As with previous pQTL studies of similar size in DO mice (Chick et al., 2016; Gyuricza et al., 2022), local pQTL tend to be more significant than distant pQTL (local median LOD = 10.8; distant median LOD = 7.9), and for over 80% of genes that have a local pQTL, we also detected an eQTL for the cognate transcript. For most of these local eQTL-pQTL pairs, the founder strain allele effects at the peak SNP are highly correlated (75% of local pairs are significant at FDR < 0.05; median *r* = 0.9), consistent with a single causal variant driving both transcript and protein abundance (Fig 4B). Correlation between chromatin accessibility (caQTL) and co-mapping local pQTL is more variable, with some proximal caQTL showing strong correlation of allele effects and others showing little or even negative correlation to local pQTL. For example, a chromatin region within the promoter of *Bspry*, a gene linked to pluripotency in mESCs and early embryonic development (Ikeda et al., 2012), has a local caQTL with highly concordant founder allele effects on *Bspry* transcript and protein abundance (Fig 4C, top). Anticorrelated local caQTL include a variable region in the promoter of the gene *Tfcp2l1* which encodes a transcription factor that has critical roles in maintenance of naïve pluripotency (Qiu et al., 2015; Ye et al., 2013). The founder allele effects at this caQTL are nearly opposite to those for the *Tfcp2l1* local eQTL and pQTL (*r* = −0.8 for both caQTL-eQTL and caQTL-pQTL pairs) (Fig 4C, bottom). Both strongly positively and negatively correlated local effects may still implicate a single causal variant but have different molecular mechanisms, e.g., a promoter variant bound by a repressor could explain the anticorrelated caQTL and pQTL for *Tfcp2l1*, whereas uncorrelated founder effects implicate multiple causal variants having distinct, unrelated effects on chromatin accessibility and transcript/protein abundance.

**Figure 4:**
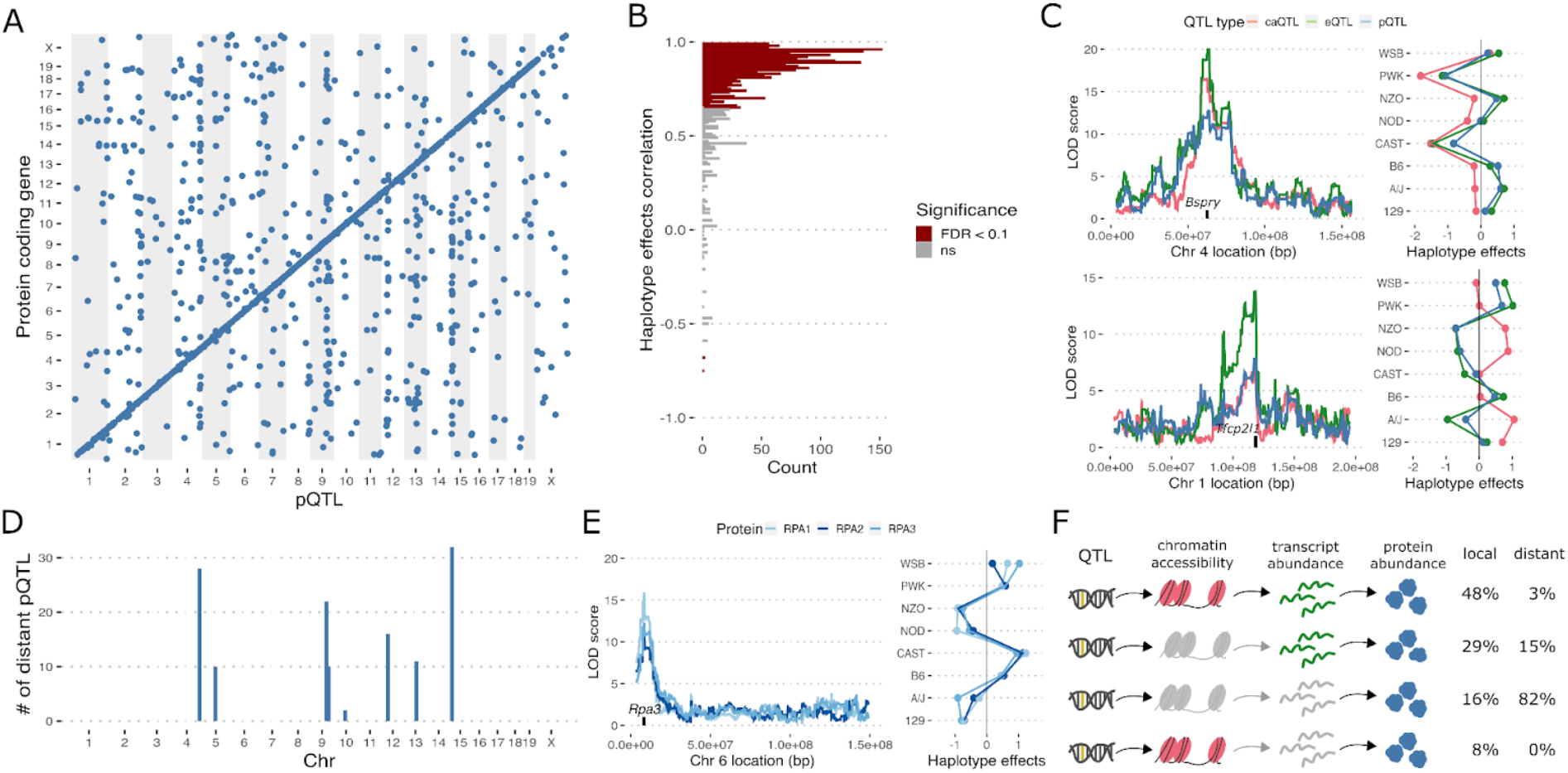
Genetic characterization of the pluripotent proteome. (A) Genetic mapping identifies 1,677 significant pQTL (LOD >7.5, permutation genome-wide p < 0.05, FDR = 0.06) where 1,056 are local (within 5Mb of the protein coding gene, seen on the diagonal) and 621 are distant (off the diagonal). pQTL are plotted across the genome where the x-axis shows the location of the pQTL and the y-axis shows the midpoint of the protein coding gene. (B) Majority of co-mapping eQTL and pQTL show high agreement in haplotype effects. Histogram of pairwise Pearson correlation coefficients between inferred allele effects from eQTL and pQTL scans for each gene with a co-mapping QTL. The bars are colored by the significance of the pairwise correlation. (C) Examples of significant pQTL where the influence of genetic variation is seen at all three molecular layers are shown. On the left, LOD scores obtained from genome scans using chromatin accessibility (caQTL), transcript (eQTL) and protein abundance (pQTL) of the associated gene is plotted with the protein coding gene location annotated on the x-axis. On the right, haplotype effects obtained from the caQTL, eQTL and pQTL peaks are shown. (D) Histogram depicting the number of total distant pQTL at hotspots across the genome. (E) An example of physical interaction propagating the effects of genetic variation is plotted. On the left, genome scan showing LOD scores across the genome for proteins RPA1, RPA2 and RPA3 is shown with the location of *Rpa3* gene annotated on the x-axis. On the right, the inferred founder allele effects at the pQTL peak for all three genes are shown. (F) Graphical overview of the different groups of pQTL where the genetic variation (QTL) influences one or more molecular layers. Molecular layers lacking any impact (i.e., no QTL above LOD >5 with matching haplotype effects) are depicted in gray.

Local pQTL likely reflect *cis*-regulatory or nonsynonymous coding variants, whereas distant pQTL are *trans* effects and likely mediated through another protein such as a transcription factor. Distant pQTL are not uniformly distributed across the genome and co-locate at pQTL “hotspots”, as we previously observed for caQTL and eQTL in DO mESCs (Skelly et al., 2020). We identified three pQTL hotspots on Chromosomes (Chrs) 4, 9, and 15 (Fig 4D), two of which (Chrs 4 and 15) were previously mapped as caQTL and/or eQTL (Skelly et al., 2020) while the Chr 9 hotspot uniquely affects protein levels (Table S4). The identity of the causal gene underlying the Chr 9 pQTL remains to be established, but targets of this pQTL-specific hotspot are enriched for proteins involved in translation initiation. This hotspot has not been detected in pQTL analyses of adult DO tissues, and may point to a post-transcriptional regulatory mechanism that is unique to pluripotent mESCs. By contrast, we previously discovered a caQTL-eQTL hotspot on Chr 15 with shared transcriptional effects on hundreds of transcripts and chromatin peaks; the Chr 15 pQTL hotspot maps to the same region and exhibits similar properties. Indeed, we observe the same founder allele effects and we identified *Lifr* transcript as the top candidate mediator for most pQTL that map to this locus (Fig S4A, B), consistent with previous findings for caQTL and eQTL (Skelly et al., 2020). We were unable to detect LIFR protein in our mass spectrometry data likely because it is a transmembrane protein with low solubility (Schey et al., 2013). Among the 32 significant pQTL at this hotspot, 14 are found only for proteins, including TCF7L1, a regulator of exit from pluripotency (Kalkan and Smith, 2014). While these unique pQTL could reflect post-transcriptional effects from LIFR, we find it more likely that transcript abundance for these genes is affected by variation in *Lifr* expression but the eQTL failed to reach our detection threshold. Likewise, of the 107 protein-coding genes with significant Chr 15 eQTLs we identified previously, only 9 are detected here as significant pQTLs. Again, many of these are likely false negatives due to the stringent genome-wide detection threshold. Finally, we treated our protein GSVA sample enrichment scores (Table S2) as quantitative traits for mapping, and find that the protein ADP-ribosylation pathway maps with a near significant QTL on proximal Chr 15 (LOD = 7.4, FDR = 0.06) that is best mediated by *Lifr* transcript abundance (Fig S4C), confirming and explaining its correlation to *Lifr* genotype in Figure 1D.

Physical interactions among proteins can propagate or buffer the effects of transcriptional variation on protein abundance (Chick et al., 2016; Keele et al., 2021; Mirauta et al., 2020). This “stoichiometric buffering” significantly affects proteins that bind in stable complexes and likely accounts for their increased covariation and lower heritability in DO mESCs. As a result, we map fewer pQTL for protein complex members, consistent with previous reports (Keele et al., 2021). We find abundant evidence for stoichiometric buffering of protein complexes, for example in ribosomal and chromatin remodeling complexes where subunits vary little in their protein abundance—and consequently do not map with any pQTL—despite varying considerably in their transcript abundance and mapping with many significant eQTL. In addition, we observe protein complexes that vary extensively across DO mESCs and have significant pQTL. Local genetic variation affecting a single subunit can propagate to other members of the complex. For example, the replication complex, with subunits RPA1, RPA2, and RPA3, co-map to a pQTL on Chr 6 and have concordant founder allele effects (Fig 4E). The *Rpa3* gene is located nearby, and *Rpa3* transcript levels are affected by a local eQTL that exhibits the same founder allele effects, suggesting that the causal variant acts in *cis* and influences transcript abundance of *Rpa3* and protein abundance of all three subunits. Indeed, mediation analysis identifies RPA3 protein abundance to be the best candidate mediator for the RPA1 and RPA2 pQTL. However, rather than RPA3 being an active regulator of RPA1 and RPA2, the local variant likely adversely affects its expression and causes it to be the limiting subunit of the complex, leading to stoichiometric buffering of the two other complex members.

The impacts of genetic variants on protein abundance can be broadly classified by their genomic location and whether they are most likely to affect transcriptional or post-transcriptional processes (Fig 4f). Most local pQTL appear to stem from transcriptional variants acting in *cis* to affect local chromatin accessibility and/or transcript abundance of the protein-encoding gene; 60% (n = 1,589) of all pQTL but 84% (n = 1,008) of local pQTL show similar genetic effects across all three molecular layers (caQTL + eQTL + pQTL; n = 483) or at least chromatin accessibility (caQTL + pQTL, n = 80) or transcript abundance (eQTL + pQTL; n = 288). On the other hand, pQTL that uniquely affect protein abundance are largely distant (75% of all unique pQTL are distant; n = 476) and mediation analysis suggests these *trans* effects can stem from physical interactions between binding partners and complex members. For the small number of local pQTL that affect abundance of the protein but not its cognate transcript (n = 157), these post-transcriptional effects may be due to protein-coding variants that alter translation efficiency or stability of the protein. Overall, these data demonstrate that the high variability in the proteome observed across DO mESC lines is also highly heritable, with many of the genetic variants driving protein-level differences showing concordant effects upstream on transcript abundance and even local chromatin accessibility. Moreover, protein-specific pQTL are also abundant and demonstrate the importance of post-transcriptional regulation and physical interactions among proteins of the quantitative proteome in mESCs. A list of all significant pQTL from this study can be found in Supplemental Table S5.

### Integration of the proteome with the chromatin landscape and transcriptome reveals signatures spanning multiple layers of biological regulation

The extensive co-variation observed within and among the mESC proteome, transcriptome, and chromatin accessibility, along with numerous shared QTL that appear to affect more than one of these regulatory layers, suggest the presence of one or more overarching regulatory signatures that co-vary among the genetically diverse DO mESC lines. To characterize these sources of variation more fully, we applied multi-omics factor analysis (MOFA, (Argelaguet et al., 2018)) to integrate and map our three genomic data sets onto a smaller set of latent factors— akin to principal components—that explain a significant proportion of the variation across mESC lines. For this analysis, we included a subset of the 15,000 most variable regions of open chromatin along with the complete sets of expressed transcripts (n = 14,405) and proteins (n = 7,432). We identified 23 latent factors that capture variation within and across the multi-omics data (Fig 5A) (Table S6). Several of the latent factors correlate with biological variables that we previously identified as major drivers of variation, including chromosomal sex (Factors 1, 10, 16, 18, 20; FDR < 0.05) and genotype at the *Lifr* locus (Factors 3, 8, 14, 18, 22; (Skelly et al., 2020)). Factors differ in the degree of variation they explain both within and across datasets, and seven factors capture variability spanning at least two or more layers of genomic data. For example, Factor 4 captures 5.4% of the observed variation in transcript abundance but also explains 0.33% of variation in chromatin accessibility (Fig 5A). Factor 4 combines information across hundreds of transcripts with thousands of chromatin sites (Fig S5A). Other factors capture variation across all three layers. For example, Factor 14 explains a small amount of variation for thousands of chromatin peaks (1.7%), transcripts (0.8%), and proteins (0.6%) (Fig S5A). In all, the 23 MOFA factors explain 27%, 41%, and 36% of the variation in chromatin accessibility, transcript, and protein abundance, respectively.

**Figure 5.**
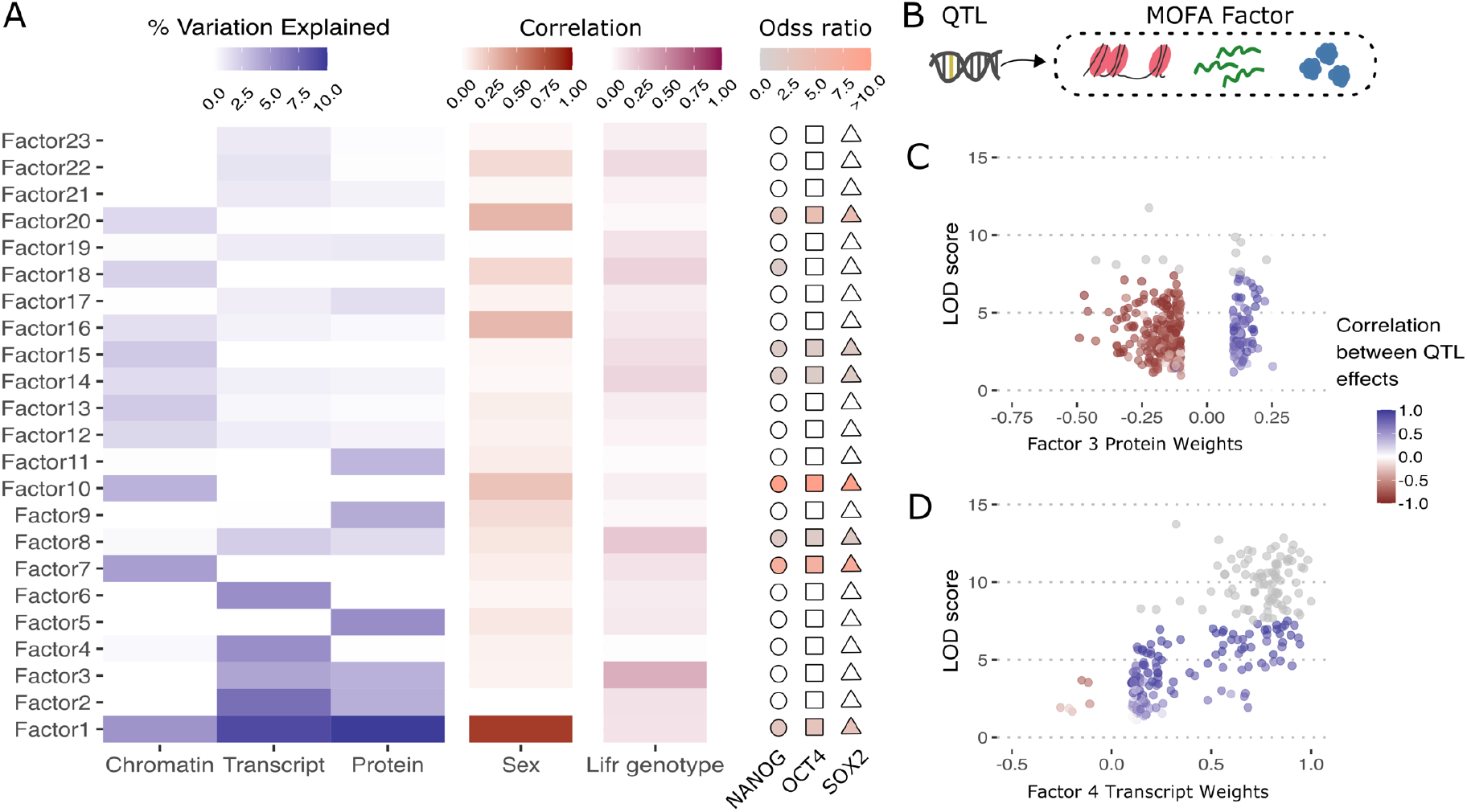
MOFA reveals broad regulatory signatures that encompass multiple layers of data. (A) MOFA yielded 23 latent factors that capture variation in one or more layers of genomic data. For each factor, percent of variation explained in chromatin accessibility, transcript abundance, and protein abundance is displayed as a heatmap, as is the correlation of each factor to experimental covariates including sex and genotype at the *Lifr* locus. Heatmap on the right indicates overrepresentation of pluripotency regulator binding sites (NANOG, OCT4 (*Pou5f1*) and SOX2) among the top chromatin drivers of each factor. (B) Depiction of QTL mapping with MOFA factors to identify the genetic modifiers driving variation across three molecular layers. (C) For all expressed proteins, the pQTL LOD score calculated at the Chr 15 QTL peak is plotted on the y-axis relative to the protein’s contribution (factor weight) to MOFA Factor 3 on the x-axis. Proteins with absolute factor weights less than 0.1 were filtered. Correlation between allele effects at the Chr 15 pQTL for individual proteins to allele effects of the Factor 3 QTL. Individual genes that mapped with a significant QTL (LOD > 7.5) are colored gray, and highlight that many proteins contribute substantially to Factor 3 and show high agreement in allele effects at the Chr 15 QTL (dark red and blue), despite not mapping individually with a significant QTL at that locus. (D) For all expressed transcripts, the eQTL LOD score at the Chr 10 QTL peak is plotted on the y-axis relative to that transcript’s contribution to Factor 4 on the x-axis. Again, transcripts with absolute factor weights less than 0.1 were filtered, and individual points are as described in panel C. Many transcripts contribute to Factor 4 and have correlated allele effects at the Chr 10 QTL, despite individually failing to map with a significant Chr 10 eQTL.

We further dissected the regulatory signatures captured by each MOFA factor through functional annotation of their molecular drivers. This included enrichment of biological processes and pathways among protein and transcript drivers ranked by factor weights, and overrepresentation of transcription factor binding sites in the genomic sequences underlying chromatin peaks. Significantly, for seven of the 23 factors, we find overrepresentation of binding sites associated with the core pluripotency factors NANOG, SOX2 and OCT4 in the sequences of their top ATAC-seq peak drivers. For three factors, including Factor 3, we find enrichment for genes involved in the regulation of pluripotency maintenance, such as response to LIF. Together, this functional evidence shows that MOFA factors are capturing variation across the molecular datasets that is relevant to pluripotency maintenance.

Twenty-two of the 23 MOFA factors have a non-zero heritability (median *h^2^* = 0.5) indicating a strong genetic contribution to their observed variability in the DO mESC lines. To identify genetic loci and causal genes driving these MOFA factors, we treated each factor as a quantitative trait and performed QTL mapping and mediation analysis (Fig 5B). We mapped 10 significant QTL across six factors (Table 1). Five of these QTL colocalize with molecular QTL hotspots described above, including Factor 3, which mapped to the *Lifr* locus (Skelly et al., 2020) (Fig 5A). The MOFA analysis identified additional transcripts and proteins that individually did not have significant association with the Chr 15 QTL but were significant contributors to Factor 3. Examination of their individual eQTL and pQTL showed evidence for subthreshold genetic association and allele effects that are consistent with regulation by the *Lifr* locus (Fig 5C). MOFA Factor 4, which captures a large amount of variation in transcript abundance, mapped to the eQTL hotspot on Chr 10. Genes mapping to this QTL include those that are upregulated in the rare 2-cell like cell state (2CLC) and are predicted to be regulated by *Duxf3* (Skelly et al., 2020). Based on their contribution to Factor 4 and shared genetic effects at the locus, we identified additional target genes known to be upregulated in the 2CLC state (n = 13) including *Zscan4e* and *Tcstv1* that individually lack significant QTL (Fig 5D) (Hendrickson et al., 2017). Mediation analysis identifies *Gm20625* transcript abundance as the best candidate regulator for this MOFA factor QTL on Chr 10 (mediation LOD drop 12 → 1.6), contradicting our previous best candidate *Duxf3* (Skelly et al., 2020), which in the current expanded analysis appears less likely to be the Chr 10 regulator (mediation LOD drop 12 → 10). Further, we identified two single nucleotide variants near *Gm20625* (rs49316493, rs265937729) that reside in annotated regulatory regions active in ESCs and that both have a founder strain genotype pattern matching the observed genetic effects at the QTL. These data implicate *Gm20625*—a gene model predicted to encode a lncRNA—as potentially playing a regulatory role in the transition between the mESC and 2CLC states. Finally, we mapped novel regulatory loci for two of the MOFA factors (Table 1), including a significant Chr 16 QTL for Factor 14 which influences hundreds of features across all three molecular layers. Mediation analysis fails to identify strong transcript or protein candidates at these novel loci, suggesting that one or more may be due to causal variants that affect the structure or function of the regulatory protein (e.g., missense variant) rather than its abundance in mESCs.

**Table 1.**
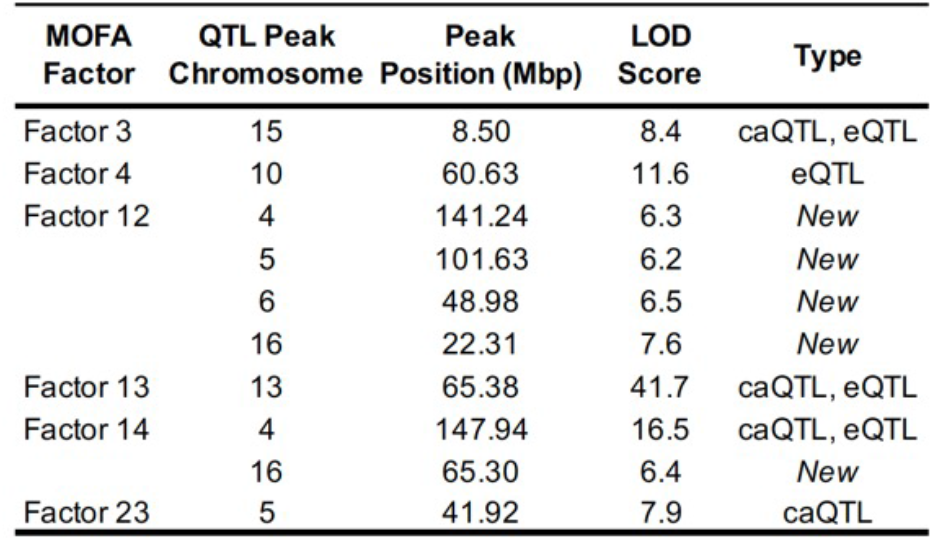
MOFA Factor QTL details for peaks above genome wide significance threshold calculated individually for each factor. Loci that were previously observed as molecular QTL hotspots are denoted in the “Type” column.

Altogether, these examples highlight the power of multi-omics data integration and factor analysis (MOFA) to reveal higher-order regulatory signatures, identify additional genes as targets (Factor 3) and mediators (Factor 4) of previously mapped QTL hotspots, and discover novel loci that influence variation across all three molecular layers (Factors 12 and 14).

## Discussion

We carried out a comprehensive genetic characterization of the pluripotent proteome in ESCs using mass spectrometry to quantify 7,432 proteins across 190 Diversity Outbred mESC lines. These data reveal that the proteome is highly variable across cell lines. Genetic background and sex are major drivers of this variation. We previously identified significant sex differences in gene expression stemming largely from X Chromosome dosage (Skelly et al., 2020), and here we find that these differences are carried through to the protein level (Schulz et al., 2014; Werner et al., 2017). Gene Set Variation Analysis (GSVA) identified multiple pluripotency and differentiation pathways that vary in activity across ESCs, including tRNA modification (Bornelöv et al., 2019); regulation of histone acetylation (Gonzales-Cope et al., 2016), intermediate filament organization (Romero et al., 2022), glutathione biosynthesis (Gu et al., 2016; Jagust et al., 2020; Xin et al., 2019), Golgi vesicle transport (Cruz et al., 2018), hippo signaling (Frum et al., 2018; Sun et al., 2020), and JUN kinase activation (Li et al., 2019) (Table S2). Of note, variation in these pathways is uniquely observed in the proteomics data.

Protein abundance is highly heritable, and we mapped pQTL for more than 20% of all detected proteins. Most of these pQTL map close to the protein-encoding gene (local pQTL) and are also detected with concordant allele effects for gene transcripts and local chromatin accessibility. We found 680 protein coding genes with significant local eQTL that do not have corresponding local pQTL even at a relaxed threshold, which could indicate buffering against transcriptional variation. Post-transcriptional buffering is most evident among the 621 distant pQTL. We found evidence for stoichiometric buffering among the members of the replication complex, where genetic variation influencing one subunit (RPA3) is propagated to other members (RPA1, RPA2). These unique distant pQTL reveal post-transcriptional genetic interactions that are not detectable in transcriptome data, adding further support to recent findings of the importance of post-transcriptional regulation in pluripotency maintenance (Chen and Hu, 2017).

Comparison of protein abundance to our earlier genetic study of transcript abundance and chromatin accessibility (Skelly et al., 2020) revealed extensive co-variation across molecular layers. We utilized multi-omics factor analysis (MOFA) to integrate the proteomics data with chromatin accessibility and transcript abundance to explore this covariation more thoroughly and summarized the three data sets into 23 latent factors. Characterization of the MOFA factors revealed shared variation in gene regulatory signatures influencing pluripotency maintenance and correlated with chromosomal sex and genotype at the *Lifr* locus. Genetic mapping and mediation of the MOFA factors identified candidate regulatory genes underlying these multi-omics signatures. We mapped QTL for MOFA factors that colocalize to both known molecular QTL hotspots and novel loci. We identified new genes as putative targets for QTL hotspots based on their significant contributions to MOFA factors and concordant allele effects between molecular and MOFA QTL. With advances in technology and decreases in cost, multi-omics profiling has emerged as a popular tool for studying gene regulation. As demonstrated here and elsewhere, integration across multiple layers of genomic data can increase our power to detect regulatory signatures underlying cell state and developmental progression (Ma et al., 2020).

Our study revealed protein variation in several known regulators of pluripotency and lineage differentiation, underscoring the variability of the pluripotent state across these genetically diverse mESC lines that may span cell states ranging from totipotent 2C-like cells to those that are poised for differentiation to one or more cell lineages. For example, TFCP2L1 is a well-characterized marker of naïve pluripotency with high levels of genetically driven protein variation across these cell lines. Previous work has suggested that differences in differentiation capacity and developmental progression can originate directly at the naïve state (Ortmann et al., 2020). How the genetic variation and variable gene regulatory states observed among DO mESCs affect their ability to differentiate into various cell lineages remains largely unexplored. Future studies will seek to characterize if and how these molecular QTL in mESCs act to bias cell fate decisions or transcriptional regulation in downstream cell lineages. Analysis with bulk molecular measurements has yielded important biological insights, however small differences in the relative proportions of specific cell types (e.g., 2C-like cells) among mESC lines may be obscured in bulk data. Future studies using single cell genomics methods will be required to measure the extent to which cellular heterogeneity contributes to the phenotypic variability observed across genetically diverse ESCs. While single cell transcriptomics and chromatin profiling are now reasonably mature technologies, our study indicates that the picture may remain incomplete without the addition of single cell proteomics data.

## Acknowledgments

We thank Ann Wells, Greg Carter, and Martin Pera for helpful discussions and feedback on the project and manuscript. Funding sources included: NIH grants R35GM133495 to S.C.M.; R01GM070683 to G.A.C.; R35GM133724 and T32HD007065 to C.L.B.; OD010921 and OD011102 to L.G.R.; NIEHS-National Toxicology Program (NTP) HHSN273201500196P to T.C.; F32GM134599 to G.R.K.; R01GM067945 to S.P.G.; and The Jackson Laboratory (to S.C.M., C.L.B., L.G.R., G.A.C.).

## Author Contributions

Conceptualization: S.C.M., S.A., G.A.C., C.L.B., L.G.R.

Methodology: S.A., S.C.M., G.A.C., C.L.B., L.G.R., D.A.S., S.P.G., T.Z., G.R.K., D.T.P.

Investigation: S.A., D.T.P., T.Z., M.P.

Data Curation & Formal Analysis: S.A., D.T.P, S.C.M., G.A.C.

Visualization: S.A., S.C.M., D.A.S.

Writing - Original Draft, Review, & Editing: S.A., S.C.M., G.A.C., C.L.B., L.G.R., S.P.G, T.Z., D.A.S., T.C.

Supervision: S.C.M., G.A.C., C.L.B., L.G.R., S.P.G.

Project Administration: S.C.M., G.A.C., C.L.B., L.G.R.

Funding Acquisition: S.C.M., G.A.C., C.L.B., L.G.R., T.C., S.P.G.

## Methods

### Diversity Outbred mESC lines

Mouse embryonic stem cell lines were derived from male and female Diversity Outbred mice (JR #009376, The Jackson Laboratory) and maintained at Predictive Biology, Inc. as previously described (Skelly et al., 2020). For proteomics analysis, ~100,000 cryopreserved DO mESCs from each line were sent from Predictive Biology to the Gygi Lab at Harvard Medical School.

### Diversity Outbred mESC RNA-seq

Raw RNA-seq data was retrieved (ArrayExpress: E-MTAB-7728) and analyzed as previously described (Skelly et al., 2020), but using both paired-end sequencing reads instead of single end. Briefly, we aligned paired-end 75 bp reads with bowtie v1.1.2 (Langmead et al., 2009) to a pooled “8-way” transcriptome containing strain-specific isoform sequences from all eight DO founder strains, then resolved multi-mapping reads and estimated transcript- and gene-level abundance for each sample using the EMASE method as implemented in gbrs v0.1.6 (Choi et al., 2020; Raghupathy et al., 2018). Genes with a median TPM (transcripts per million) value smaller than 0.5 or zero value (i.e., not expressed) in more than half of the samples were filtered. Next, we normalized gene-level counts to the upper quartile value to account for differences in library size and then applied the ComBAT function from R/sva package to remove batch effects caused by library preparation (Johnson et al., 2007). For QTL mapping, we transformed normalized values to rank normal scores using rankZ normalization as implemented in the DOQTL R package (Gatti et al., 2014). Finally, sample mix-ups were resolved by comparing the genotypes inferred from the RNA-seq data using gbrs v0.1.6 (http://churchill-lab.github.io/gbrs/) to genotypes inferred from DNA microarrays (GigaMUGA platform, Neogen Geneseek).

### Diversity Outbred mESC ATAC-seq

Normalized ATAC-seq peak values from Skelly et al., (2020) were further processed using the ComBAT function in the R/sva package to remove any potential batch effects caused by library preparation (Johnson et al., 2007). Normalized, batch-corrected peak values were used in all correlation analyses. For QTL mapping, these values were further transformed to rank normal scores using the rankZ function from the DOQTL package (Gatti et al., 2014). For annotation of ATAC-seq peaks we utilized the ChIPseeker R package (Yu et al., 2015).

### Multiplexed quantitative proteomics analysis of DO mESCs

#### Sample preparation for proteomics analysis

Frozen cell pellets were resuspended in 8 M Urea, 200 mM EPPS, pH 8.5, with protease inhibitor, and lysed by passing through a 21-gauge needle with syringe. After centrifugation at 13,000 rpm at 4°C for 10min, supernatant was used for further analysis. BCA assay was performed to determine protein concentration of each sample. Samples were reduced in 5 mM TCEP for 15min, alkylated with 10 mM iodoacetamide for 15min, and quenched with 15 mM DTT for 15min. 200 *μ*g protein was chloroform-methanol precipitated and re-suspended in 200 *μ*L 200 mM EPPS (pH 8.5). Protein was digested by Lys-C at a 1:100 protease-to-peptide ratio overnight at room temperature with gentle shaking. Trypsin was used for further digestion for 6 hours at 37°C at 1:100. 100 *μ*L of each sample were aliquoted. 30 *μ*L acetonitrile (ACN) was added into each sample to 30% final volume. 200 *μ*g TMT reagent (126, 127N, 127C, 128N, 128C, 129N, 129C, 130N, 130C, 131N) in 10 *μ*L ACN was added to each sample. After 1 hour of labeling, 2 *μ*L of each sample was combined, desalted, and analyzed using mass spectrometry. TMT labeling efficiency was calculated and over 99%. After quenching using 0.3% hydroxylamine, 10 samples in each TMT were combined and fractionated with basic pH reversed phase (BPRP) high performance liquid chromatography (HPLC), collected onto a 96 six well plate and combined for 24 fractions in total. Twelve fractions were desalted and analyzed by liquid chromatography-tandem mass spectrometry (LC-MS/MS) (Navarrete-Perea et al., 2018).

#### Liquid chromatography and tandem mass spectrometry

For the BPRP fractions, mass spectrometric data were collected on an Orbitrap Fusion mass spectrometer coupled to a Proxeon NanoLC-1200 UHPLC. The 100 *μ*m capillary column was packed with 35 cm of Accucore 50 resin (2.6 μm, 150Å; ThermoFisher Scientific). The mobile phase was 5% acetonitrile, 0.125% formic acid (A) and 95% acetonitrile, 0.125% formic acid (B). The data were collected using a DDA-SPS-MS3 method. Each fraction was eluted using a 150 min method over a gradient from 6% to 30% B. Peptides were ionized with a spray voltage of 2,600 kV. The instrument method included Orbitrap MS1 scans (resolution of 1.2 x105; mass range 350-1400 m/z; automatic gain control (AGC) target 5×105, max injection time of 100 ms and ion trap MS2 scans (CID collision energy of 35%; AGC target 2×104; rapid scan mode; max injection time of 120 ms). MS3 precursors were fragmented by HCD and analyzed using the Orbitrap (NCE 65%, AGC 1 x105, maximum injection time 150 ms, resolution was 5 x104 at 400 Th). Detailed parameters for MS2 and MS3 are embedded in the RAW files.

#### Mass spectrometry data analysis

Mass spectra were processed using a Sequest-based pipeline (Huttlin et al., 2010). Spectra were converted to mzXML using a modified version of ReAdW.exe. Database search included all entries from an indexed Ensembl database version 90 (downloaded:10/09/2017). This database was concatenated with one composed of all protein sequences in the reversed order. Searches were performed using a 50 ppm precursor ion tolerance for total protein level analysis. The product ion tolerance was set to 0.9 Da. TMT tags on lysine residues and peptide N termini (+229.163 Da) and carbamidomethylation of cysteine residues (+57.021 Da) were set as static modifications, while oxidation of methionine residues (+15.995 Da) was set as a variable modification. In addition, for phosphopeptide analysis, phosphorylation (+79.966 Da) on serine, threonine, and tyrosine are included as variable modifications. Peptide-spectrum matches (PSMs) were adjusted to a 1% false discovery rate (FDR). PSM filtering was performed using a linear discriminant analysis (LDA). For TMT-based reporter ion quantitation, we extracted the summed signal-to-noise (S:N) ratio for each TMT channel and found the closest matching centroid to the expected mass of the TMT reporter ion. For protein-level comparisons, PSMs were identified, quantified, and collapsed to a 1% peptide false discovery rate (FDR) and then collapsed further to a final protein-level FDR of 1%, which resulted in a final peptide level FDR of <0.1%. Moreover, protein assembly was guided by principles of parsimony to produce the smallest set of proteins necessary to account for all observed peptides. Proteins were quantified by summing reporter ion counts across all matching PSMs. PSMs with poor quality, MS3 spectra with less than 10 TMT reporter ion channels missing, MS3 spectra with TMT reporter summed signal-to-noise of less than 100 or having no MS3 spectra were excluded from quantification. Each reporter ion channel was summed across all quantified proteins and normalized assuming equal protein loading of all 10 samples. The mass spectrometry proteomics data have been deposited to the ProteomeXchange Consortium via the PRIDE partner repository with dataset identifiers PXD033001.

#### Protein abundance estimation

Protein abundances were estimated as described previously (Keele et al., 2021). Briefly, peptides that contain polymorphisms were filtered and batch effects were removed from the filtered peptide data using a linear mixed model fit with the R/lme4 package (Bates et al., 2014). Finally, protein abundances were estimated and normalized using the processed peptide data as described in detail in Keele et al., (2021). Proteins missing values in more than 50% of the samples were removed from further analysis.

### Statistical Analysis and Genetic Mapping

#### Code availability

All analyses and figures were generated with the R statistical programming language and are available at the following web resource (link here) and github (link here). Unless otherwise stated R/tidyr package was used for data processing, R/ggplot2 for plotting and R/pheatmap for heatmap plots.

#### Gene annotations and id matching across data sets

Transcript abundance data was annotated to Ensembl gene identifiers, proteomics data was annotated to Ensembl protein identifiers, and ATAC-seq data was annotated to Ensembl gene ids using ChipSeeker R package. We used ENSEMBL v98 to add gene annotations such as MGI symbol, gene location, and gene biotype. MGI symbol was used as the identifier for all downstream analysis such as overrepresentation and gene set enrichment.

#### Correlation Analysis

We used the rcorr function from the R/Hmisc package to calculate Pearson correlations. Individual p-values were adjusted for multiple testing using the p.adjust function in R/base and specifying the Benjamini-Hochberg (“BH”) option to estimate the false discovery rate (FDR).

##### Sample-to-sample correlation for protein abundance

For proteome-to-proteome comparisons, we used the abundance of 7,432 proteins across 190 cell lines. To compare chromatin accessibility profiles to the proteome, we used 36,859 ATAC-seq peaks annotated to 6,865 proteins and their corresponding protein abundances in 163 cell lines for which ATAC-seq, transcriptomics and proteomics were profiled. Similarly, for comparing the transcriptome to the proteome across 174 cell lines that had both RNA-seq and proteomics data, we used the overlapping set of 7,241 genes with both transcript and protein abundance measures.

##### Correlation between chromatin accessibility and protein abundance

Pairwise Pearson correlations were calculated between the abundance of 7,148 autosomal proteins and the chromatin accessibility of 99,159 autosomal ATAC-seq peaks across 163 cell lines for which ATAC-seq, transcriptomics and proteomics were profiled.

##### Correlation between transcript and protein abundance for individual genes

Pairwise Pearson correlations were calculated for 7,241 genes with both transcript and protein abundance measures across 174 cell lines that had both RNA-seq and proteomics data.

##### Correlation between complex member and non-complex member proteins

The list of complex member proteins was retrieved from (Romanov et al., 2019) which includes protein complexes manually curated using CORUM and COMPLEAT databases. Pairwise Pearson correlations between protein abundances of complex member and non-complex member genes were calculated for complexes with five or more subunits (n = 164) excluding proteins with significant pQTL to leave out large genetic effects that may not be shared among complex members.

#### Gene Set Enrichment and Over-representation Analysis

We performed over-representation analysis using the ‘gost’ function in the gProfiler2 package in R (Raudvere et al., 2019) using an appropriate universal background on a case-by-case basis and ‘fdr’ option for p-value correction. For example, when looking at the functional enrichments in proteins with high variation all genes identified in proteomics were used whereas only the shared set of genes between RNA-seq and proteomics was used when looking at genes with positive correlation between transcript and protein abundance. For gene set enrichment analysis, we used the WebsGestaltR R package (Liao et al., 2019). To identify overrepresentation of genomic regions we utilized R package LOLA (Sheffield and Bock, 2016) which looks at the overlap between user data sets and public genomic data sets like transcription factor binding sites from ENCODE and the CODEX database. Following instructions of the R/LOLA package the p-values were transformed to q-values using the R/qvalue package to get FDR values.

#### Gene Set Variation Analysis

We performed Gene Set Variation Analysis using the R/Bioconductor package GSVA (Hänzelmann et al., 2013). Gene Ontology terms with gene symbols were retrieved from MGI (http://www.informatics.jax.org/gotools/data/input/MGIgenes_by_GOid.txt) which included 8,436 GO Biological Process, 3,355 GO Molecular Function and 1,077 GO Cellular Component gene sets. List of protein complexes and subunits was retrieved from (Romanov et al., 2019) which includes protein complexes manually curated using CORUM and COMPLEAT databases. Enrichment scores were calculated using the abundance of 7,432 proteins across 190 cell lines for each gene set with at least 5 overlapping proteins. Next, we evaluated the significance of enrichment scores across experimental covariates using a two-way ANOVA (~ sex + *Lifr* genotype *+ sex:Lifr* genotype) where individual p-values were corrected for multiple testing using the p.adjust function in R/base and specifying the Benjamini-Hochberg (“BH”) option and followed by Tukey’s HSD using R/rstatix package for pairwise comparisons. Categories that showed significance in both statistical tests were reported with the p-value obtained from Tukey’s HSD.

#### Quantitative Trait Locus Mapping

Genetic mapping was performed using a linear-mixed model implemented as the ‘scan1’ function in R/qtl2 package (Broman et al., 2019). We mapped using the normalized, transformed values with sex as a covariate and the Leave One Chromosome Out (loco) option for kinship correction (Gatti et al., 2014). To estimate genome-wide significance, we permuted genotypes 1000 times while maintaining the relationship between the phenotype and covariates. For each permutation we retained the maximum LOD score in order to generate a null distribution for the test statistic (Churchill and Doerge, 1994). To calculate thresholds for pQTL, we repeated this permutation strategy for all proteins and estimated a significance cutoff at LOD > 7.5 (alpha = 0.05), and a suggestive cutoff at LOD > 6. False discovery rates (q-values) were determined for each permutation-derived p-value with R/Qvalue software, using the bootstrap method to estimate ⊓_0_ and the default λ tuning parameters (Storey et al., 2004). Support intervals for each QTL were defined by the 95% Bayesian credible interval (Sen and Churchill, 2001). We call a QTL ‘local’ if the QTL peak is within 5Mbp to the midpoint of its corresponding gene and ‘distal’ if otherwise. Founder allele effects were estimated as best linear unbiased predictors (BLUPs) at the QTL using scan1blup function in R/qtl2 package. Previous work has estimated the genome-wide significance threshold at 7.6 and 7.5 for chromatin accessibility QTL (caQTL) and expression QTL (eQTL) respectively (Skelly et al., 2020). To identify overlaps with significant pQTL, we used a relaxed threshold of LOD > 5 for caQTL and eQTL. They were classified as shared if the QTL peaks were within +/-5Mb of the significant pQTL peak and the absolute correlation between haplotype effects was higher than 0.5.

#### Defining QTL Hotspots

We first identified distal QTL that reach genome-wide permutation-based threshold (p < 0.05; LOD 7.5). Next, we applied a sliding window method to identify hotspots as described in Skelly et al., 2020. Briefly, we counted the number of distal QTL within 1cM windows (0.25 cM shift) across the genome and selected the top 0.5% of bins with the most distant pQTL (0.5% bin threshold ≥ 8 distant pQTLs). Final coordinates for each hotspot were determined using the Bioconductor package ‘GenomicRanges’ to merge adjacent bins into a single region (Lawrence et al., 2013).

#### Mediation Analysis

We used mediation analysis to identify regions of open chromatin, transcript, and protein abundance that were likely to be the causal mediator of a caQTL, eQTL, or pQTL. Mediation analysis was performed using the ‘intermediate’ package in R (https://github.com/simecek/intermediate) by regressing each target (T) on a mediator (M) at the QTL (Q) and adjusting for covariates. We applied the ‘double-lod-diff’ method to reduce the effects of missing values. For mediation of QTL with the matching data type we used the full sample set, e.g., pQTL mediation by proteins (Q_pQTL_ → Protein_M_ → Protein_⊤_) were done using all the 190 samples. On the other hand, mediation across data types were done on common set of samples e.g., for mediation between protein and transcript (Q_pQTL_ → Transcript_M_ → Protein_⊤_ | Q_eQTL_ → Protein_M_ → Transcript_⊤_ only the 174 samples with both protein and transcript measurements were used. To assess the significance of a LOD drop, we mediated the QTL against all of the mediator data, converted the recorded LOD scores to normal scores, and checked if the score fell below 6 standard deviations from the mean (Chick et al., 2016). Mediators were further filtered to narrow down top candidates to include genes with midpoints that are found within 5Mb of the QTL peak.

#### Data Integration and Multi-Omics Factor Analysis

For data integration we used Multi-Omics Factor Analysis (MOFA) implemented in Python (mofapy2) and in R (MOFA2) (Argelaguet et al., 2018). MOFA integrates multi-omics data sets in an unsupervised fashion using a factor analysis model and infers a number of interpretable latent factors. All transcripts (n = 14,405), proteins (n = 7,432) and the most variable 15,000 ATAC-seq peaks based on total variance were used for integration from 163 cell lines with all three molecular measurements. All three datasets were log transformed using base R function log1p before modeling with MOFA. For model generation, we modified the following options from default: we set number of factors to 30, number of maximum iterations to 10,000, convergence mode to “slow” and scale views option to TRUE. The model with the best convergence based on the evidence lower bound statistic (ELBO) was saved for further analysis. Next, factors that showed a significant correlation to the total number of expressed features and that didn’t explain more than 1% variation in at least one data set were removed resulting in 23 latent factors. We calculated the proportion of variance explained by factor per data set and the correlation between factors and experimental covariates using built-in functions in the MOFA2 R package. Functional characterization of MOFA Factors was done using the R/LOLA package for top ATAC-seq peak drivers and the R/WebsGestaltR package for transcripts and proteins. Top ATAC-seq drivers were obtained using the base R boxplot.stats function where the outliers correspond to data points that lie outside 1.5 times the interquartile range. MOFA factor weights were used to rank genes in enrichment analysis for transcripts and proteins. QTL mapping, mediation and permutation analysis with factors were done as described above using genotype probabilities from the 163 samples used in MOFA.

## Supplemental Tables

**Table S1. Over-represented annotations among lists of detected and undetected proteins.**

**Table S2. Results from Gene Set Variation Analysis.**

**Table S3. List of proteins with correlation to ATAC-seq peaks.**

**Table S4. List of pQTL hotspots and their target proteins.**

**Table S5. List of all significant pQTL in Diversity Outbred mESCs.**

**Table S6. Lists of molecular features and their weights for each of 23 MOFA factors.**

## Supplemental Figures

**Figure S1.**
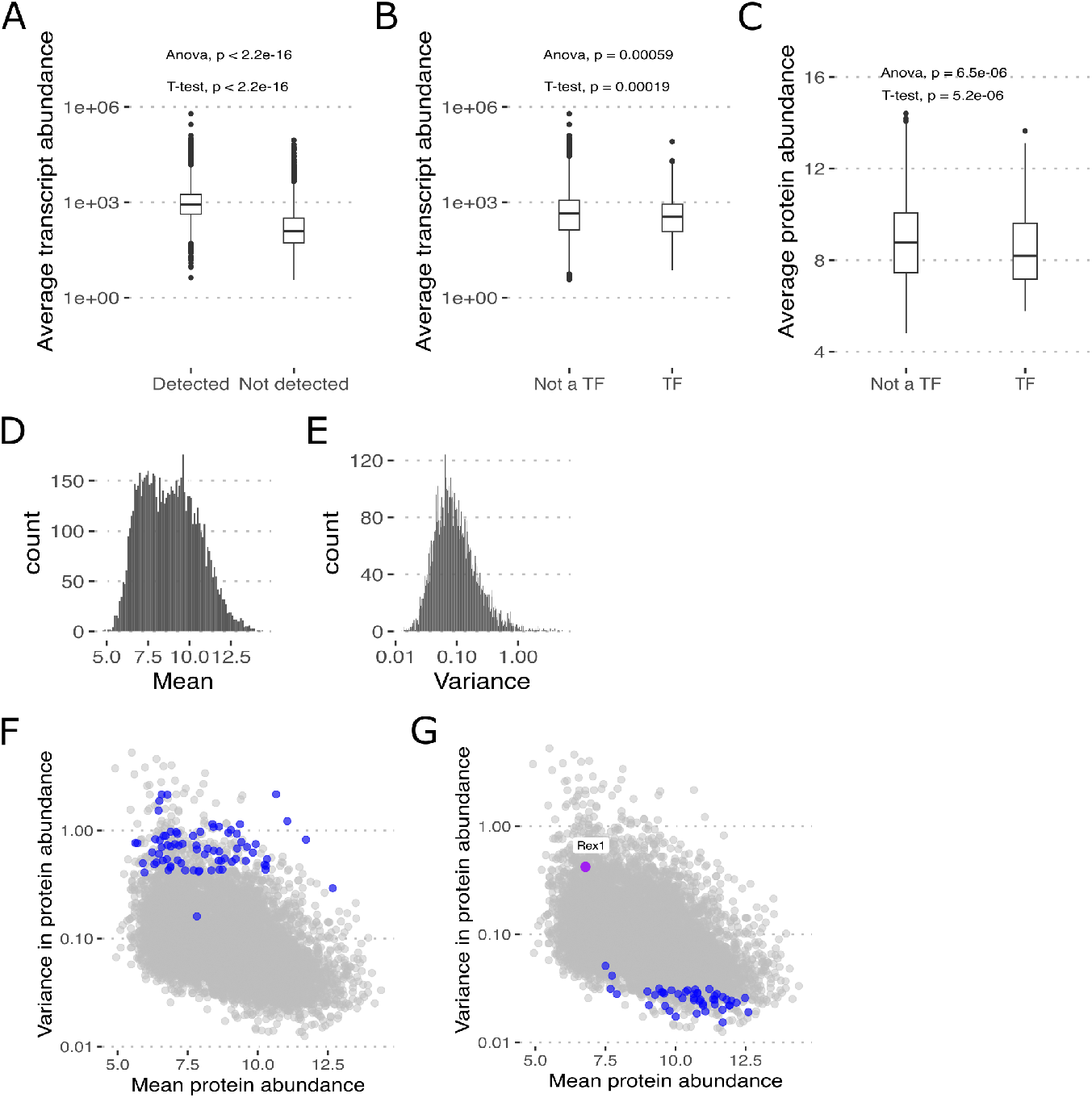
(A) Genes where protein abundance is detected have a significantly higher mean transcript abundance (One way ANOVA, followed by t-test). Average transcript abundance of protein coding genes (n = 12,732) that are detected (TRUE, n = 7,240) and not detected (FALSE, n = 5,492) in the proteomics data are plotted. (B, C) TFs show a significantly lower mean for both transcript and protein abundance in comparison to other genes (One way ANOVA, followed by t-test). Average transcript and protein abundance of protein coding genes that are transcription factors (TF) and not transcription factors (Not a TF) are plotted. (D, E) Protein abundance is highly variable across DO mESCs. Histograms showing the mean abundance and variance per protein plotted for 7,342 proteins across 190 DO mESC lines. (F) Mean abundance and variance plotted for all proteins (gray) with proteins identified as part of ‘Extracellular region’ and ‘ECM protein’ GO Terms in most variable proteins (top 5^th^ percentile %CV), in overrepresentation analysis, highlighted in blue. (G) Mean abundance and variance plotted for all proteins with proteins identified as ‘REX1 Target’ in TRANSFAC database in least variable proteins (bottom 5^th^ percentile %CV), in overrepresentation analysis, highlighted in blue and REX1 (*Zfp42*) highlighted in purple.

**Figure S2.**
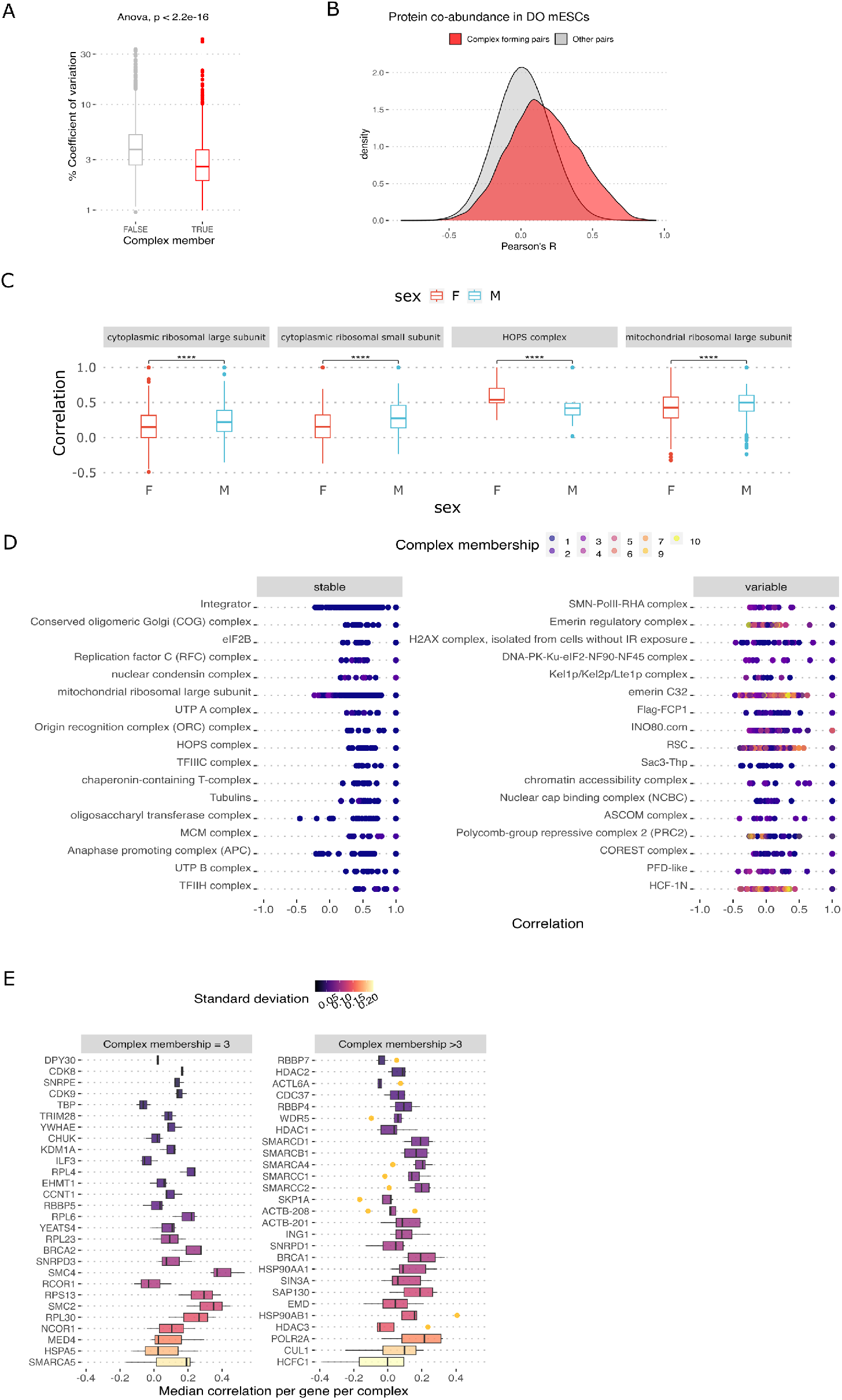
(A) Proteins that are part of a complex show less variation. Boxplots depicting % coefficient of variation of protein abundance for genes that are complex members and not complex members. (B) Proteins that physically interact show higher pairwise correlation in abundance than non-interacting proteins. Density distributions of pairwise Pearson correlations between complex forming proteins and others are plotted. (C) Sex influences the co-regulation of complex subunits. Boxplots of pairwise Pearson correlations among complex subunits with significant differences between male and female cell lines are shown (One way ANOVA followed by Tukey’s HSD, ****: p value < 0.00005) (D) Variable complexes are more likely to have promiscuous subunits that are part of more than 2 complexes. Pairwise Pearson correlation coefficients plotted for all subunits that are part of stable (upper 10th percentile, most cohesive) and variable (lower 10th percentile, least cohesive) complexes where the proteins are colored by the number of complexes they belong to. (E) Promiscuous proteins vary in preference of complexes. Boxplots of median pairwise Pearson correlations of complex subunits across various complexes they are part of are plotted. The complex subunits are separated into two categories based on the total number of complexes they belong to.

**Figure S3.**
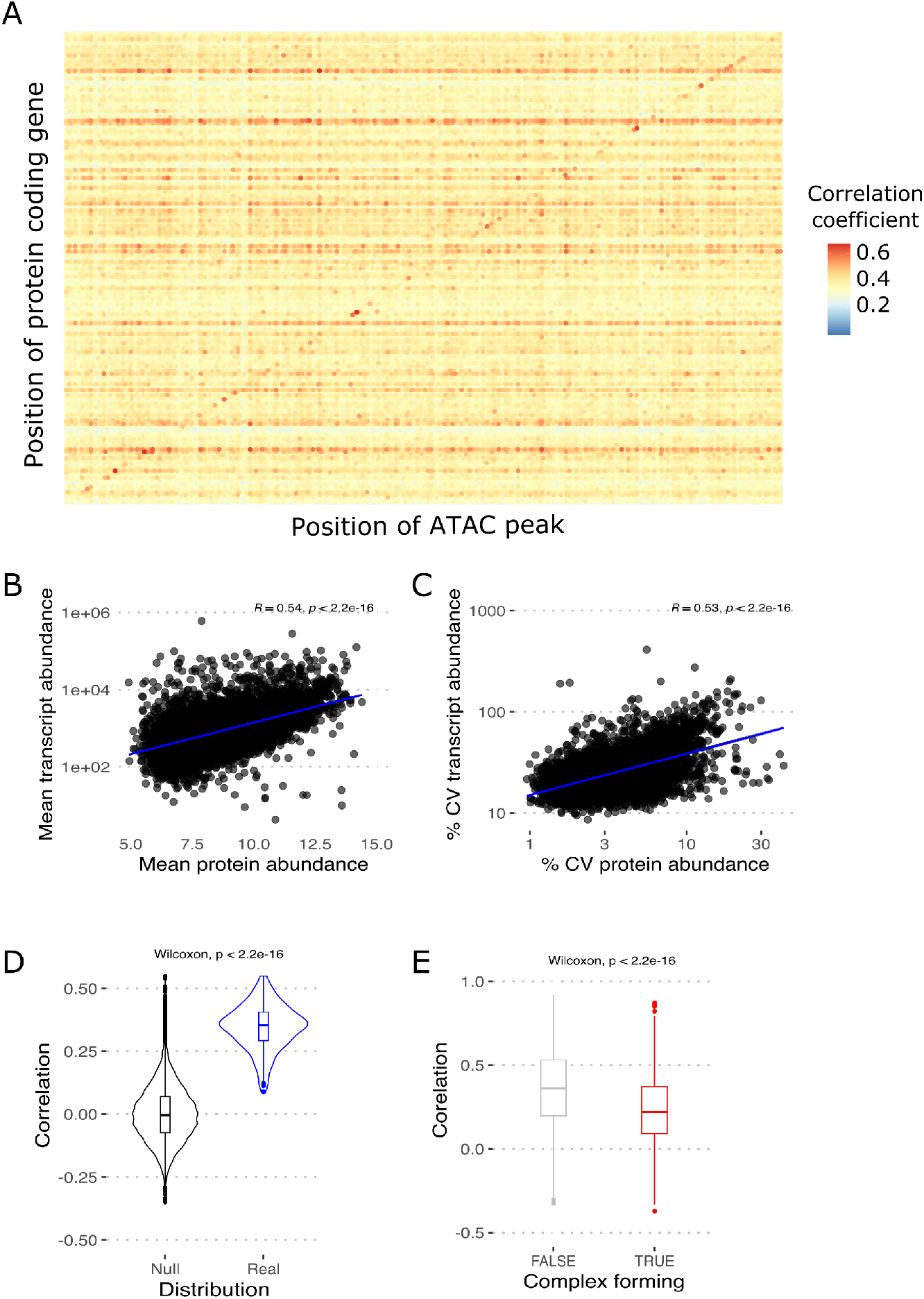
(A) A heatmap of Pearson correlation coefficients between protein abundance and chromatin accessibility across the genome. Proteins encoded on the sex chromosomes were excluded from the analysis to limit sex effects due to X gene dosage. Correlation between all autosomal proteins and accessibility at ATAC-seq peaks were calculated. For plotting, proteins and chromatin regions are grouped in 5 Kb bins and the points are colored and sized by the maximum correlation value in each bin. (B, C) Scatterplots showing mean and coefficient of variation (% CV) for transcript and protein abundance for genes with both measurements (n = 7,241). (D) Genetically identical cell lines show significantly higher correlation than what is expected by chance between the transcriptome and proteome. Violin plots overlaid with boxplots depicting the distribution of Pearson correlation coefficients between the transcriptome and proteome of genetically identical mESCs (blue) and the null distribution generated through 1000 permutations where the sample names are randomized (black). (E) Genes that do not form complexes show significantly higher correlation between transcript and protein abundance. Boxplot comparing the pairwise Pearson correlation coefficients between transcript and protein abundance for genes that are part of protein complexes (TRUE) and that do not form complexes (FALSE).

**Figure S4.**
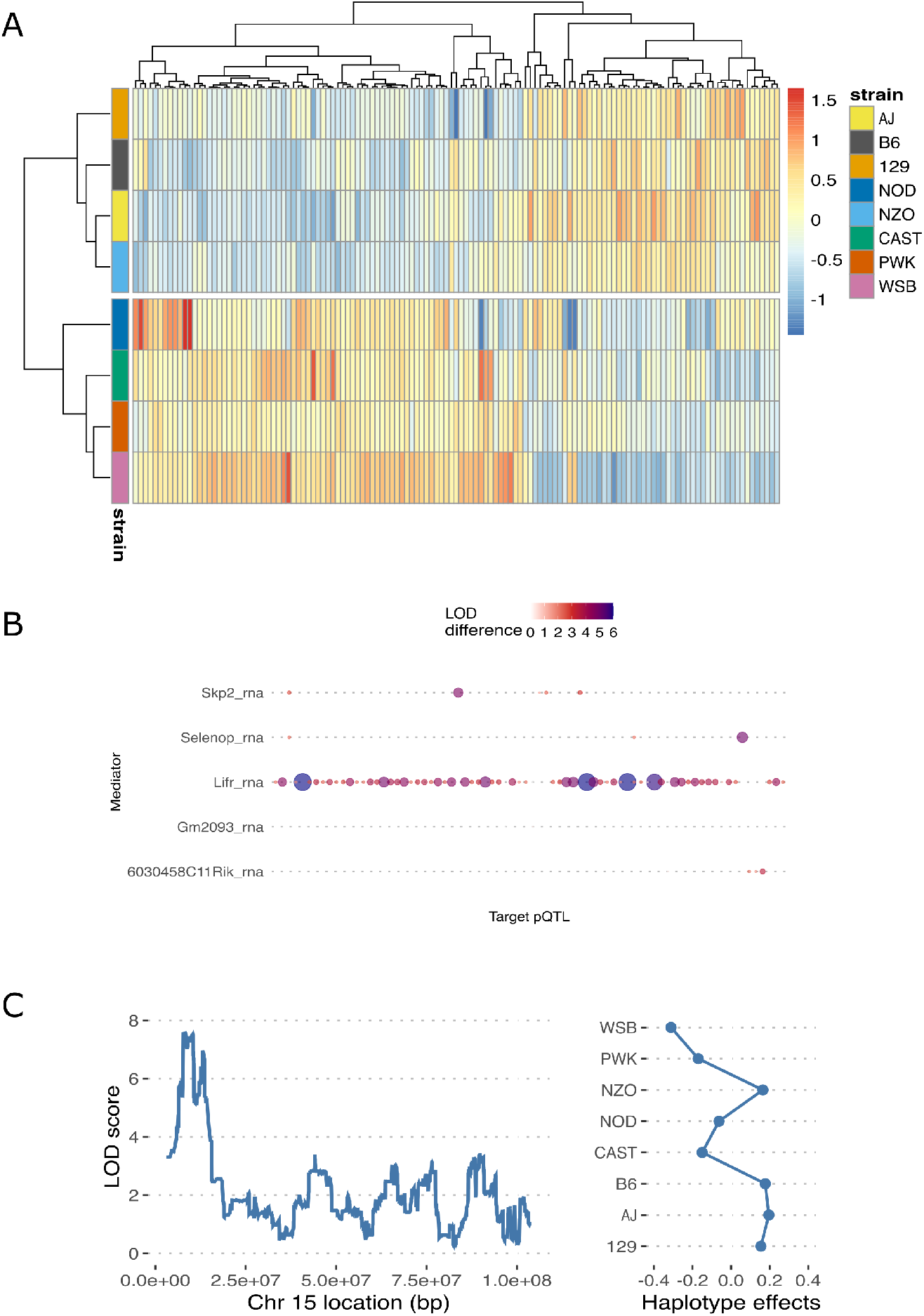
(A) The allelic split observed in previously described eQTL hotspot on chromosome 15 is also observed for the pQTL hotspot. Heatmap showing haplotype effects at suggestive distant pQTL peaks (LOD > 6) within the chromosome 15 hotspot. (B) Mediation analysis identifies *Lifr* transcript abundance as the best mediator for chromosome 15 pQTL hotspot. Decrease in LOD scores due to mediation is plotted for the top five mediators in the region for the suggestive distant pQTL. The points are colored and sized according to LOD difference. For 61/131 suggestive distant pQTL peaks in the region, *Lifr* transcript abundance leads to the largest drop in LOD when included as a covariate in the genetic mapping model. (C) Genetic mapping with GSVA scores of GO term Protein ADP-Ribosylation identifies a near significant QTL on chromosome 15 with similar haplotype effects to the chromosome 15 molecular QTL hotspot. On the left, genome scan showing LOD scores is plotted for chromosome 15. On the right, inferred haplotype effects at the QTL peak is plotted.

**Figure S5.**
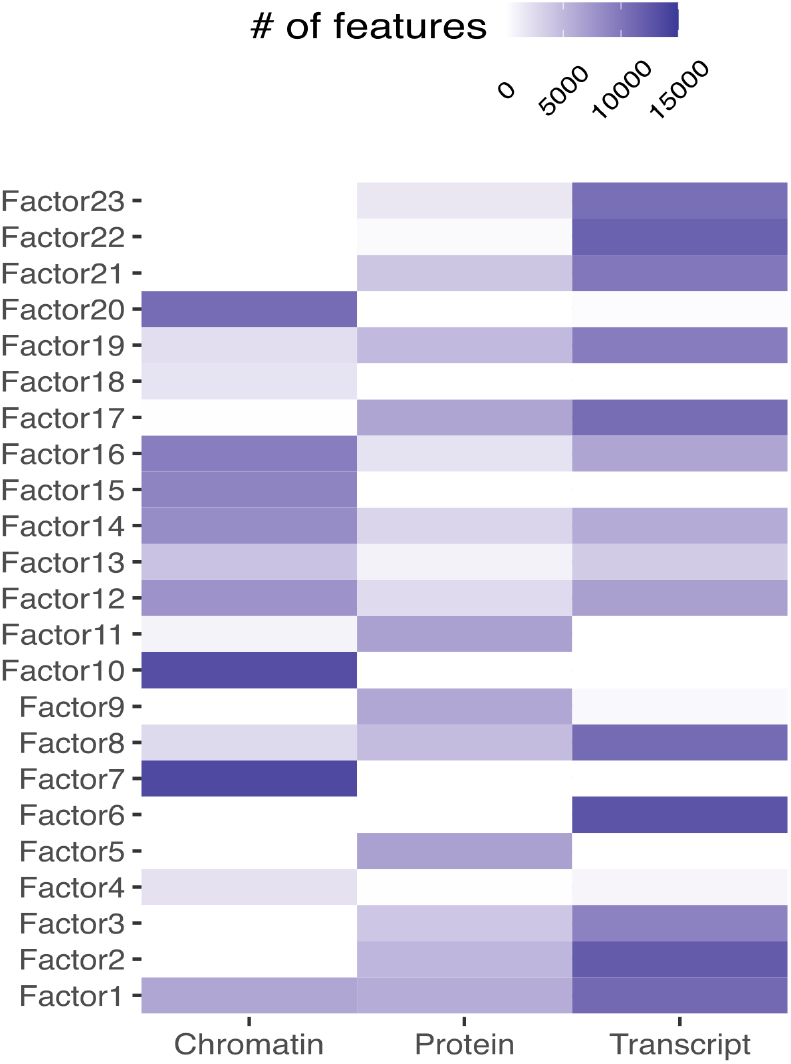
Heatmaps showing the number of features in each data set with abs(weight) > 0.01 for 23 MOFA Factors.

